# Chromosome breaks in breast cancers occur near herpes tumor virus sequences and explain why the cancer comes back

**DOI:** 10.1101/2021.11.08.467751

**Authors:** Bernard Friedenson

## Abstract

Breast cancer has a relentless tendency to come back after treatment. Analyses of public data from about 2100 breast cancers produce a model that explains this recurrence and implicates variants of Epstein-Barr viruses (EBV or Human Herpes Virus 4). These viruses cause chromosome breaks. Broken chromosome pieces rejoin abnormally, sometimes including two centromeres. Two centromeres on the same chromosome interfere with cell division. Each centromere gets pulled toward a different pole. This mechanical stress shatters chromosomes. Shattered chromosome fragments rejoin arbitrarily, but showers of mutations accompany their rejoining. In this way, a single break can destabilize the entire genome. The breast cancer phenotype is not fixed and constantly creates new cancer driver genes. The phenotype becomes independent of the original virus and its dosage. Cancer comes back because treatment does not explicitly target the underlying breakage-rejoining cycles or the contributing virus.

The following data support this model. EBV causes chromosome breaks, and breast cancer chromosomes often have two centromeres. Breast cancer breakpoints on all chromosomes aggregate around the same positions as breakpoints in cancers definitively associated with EBV infection (nasopharyngeal cancer and endemic Burkitt’s lymphoma). Rejoined boundaries of highly fragmented chromosomes characteristic of breakage fusion cycles cluster around viral sequences. There is presumptive evidence of past infection. Human EBV sequences distribute like retrovirus transposons near dense piRNA clusters at a critical MHC-immune response region of chromosome 6. Other viruses strongly resemble endogenous transposons which piRNAs inactivate by methylation and cleavage. Remnants of exogenous EBV variants sit close to inactive transposons in piRNA sandwiches. The arrangement grossly resembles bacterial CRISPR and adds a layer of DNA protection to the immune system. Breast cancers target this protection with chromosome breaks and mutations and have a distinctive methylation signature nearby. Finally, areas near EBV docking sites can have increased numbers of breaks.

**Graphical Abstract:** 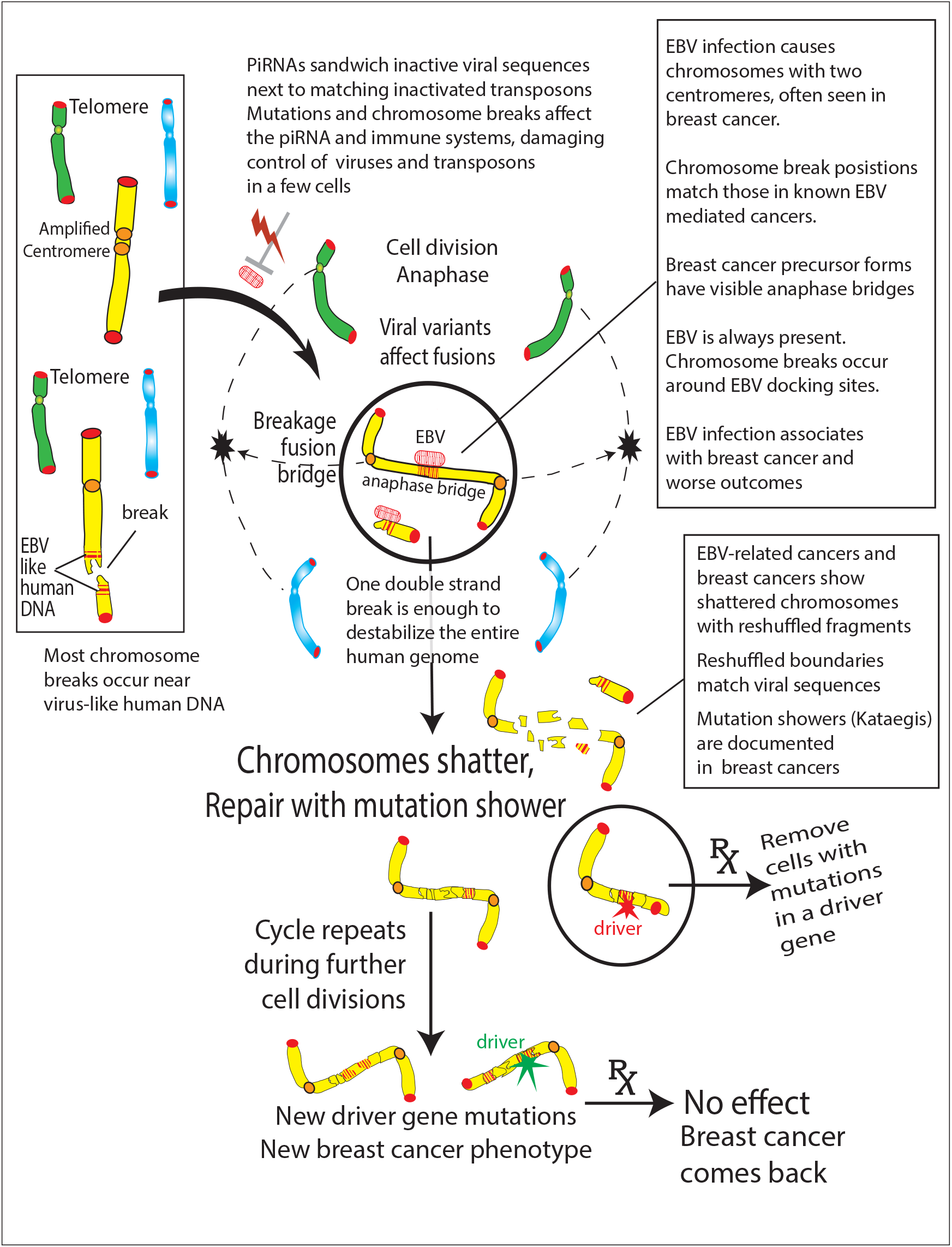

## Introduction

Breast cancer often comes back after therapy with unrelenting recurrence rates over at least 20 years. Depending on the number of lymph nodes involved, recurrence rates range from 12% (0 nodes) to 41% (4-9 nodes) (Pan *et al*, 2017). One explanation for this recurrence is that treatment has overlooked an underlying cause of the disease. By the time breast cancer is diagnosed, its causes are difficult to isolate from multiple risk factors, heredity, thousands of mutations, and a lifetime of exposure to mutagens and infections.

Human cancer viruses represent risk factors that usually just initiate or promote cancer and do not cause cancer by themselves (Sarid and Gao, 2011). For example, Epstein-Barr virus /Human herpesvirus 4 (EBV/HHV4) exists as a latent infection in almost all humans (>90%). Infection occurs early, with half the children already seropositive by age 6-8. At age 18-19, 89% of the population is seropositive (Balfour *et al*, 2013). Most people control EBV infection (Fina *et al*, 2001; Lawson and Glenn, 2021), Life-long infections of EBV in most people obscure potential associations with breast cancer, but some associations are undeniable. EBV increases breast cancer risk by 4.75 to 6.29-fold according to meta-analyses of 16 or 10 studies, respectively. An analysis of 24 case-control studies showed EBV is significantly more prevalent in breast cancer tissues than in normal and benign controls (Lawson and Glenn, 2021).

In breast epithelial cell models, EBV infection facilitates malignant transformation and tumor formation (Hu *et al*, 2016). Breast cancer cells from biopsies of sporadic and hereditary breast cancers express latent EBV infection gene products (Ayee *et al*, 2020; Lawson and Glenn, 2021), even after excluding the possibility that the virus comes from lymphocytes (Lorenzetti *et al*, 2010). Finding the replicative form of EBV in breast cancer correlates with poorer outcomes. Latent infection confers a survival advantage and accompanies anti-tumoral immune responses (Marrao *et al*, 2014). EBV activation from its latent state causes massive changes in host chromatin methylation and structure (Buschle *et al*, 2021; Kim *et al*, 2020; Tang *et al*, 2012). Aberrant methylation in breast cancer occurs at hundreds of host gene promoters and distal sequences (Batra *et al*, 2021).

To investigate the origins of breast cancer, the present study starts with nasopharyngeal cancer (NPC) as a first model for an EBV-associated epithelial cell cancer. NPC has an explicit EBV connection, with 100% of malignant cells being EBV-positive (Germini *et al*, 2020; Hau *et al*, 2020; Xu *et al*, 2019). Nearly 8500 EBV forms exist in NPC patients with over 2100 variants in a single host, each differing only slightly from a reference viral genome (Xu *et al*, 2019). Two EBV tumor virus variants (HKHD40 and HKNPC60) are strongly associated with NPC (Hui *et al*, 2019), and they become major players in my study presented below.

In the host, mutations interfere with innate immunity and constitutively activate an inflammatory response (Bruce *et al*, 2021). NPC accompanies multiple genetic chromosome structural changes, including chromosomal gains or losses, homozygous deletions, and telomere shortening (Tan *et al*, 2018). High-level expression of a centromere protein predicts progression and poor survival (Liao *et al*, 2007). Endemic Burkitt’s lymphoma is another cancer caused by EBV. Chromosomes with two centromeres have been specifically investigated in Burkitt’s lymphoma and found to be significantly increased (Kamranvar *et al*, 2007). Other chromosome structural changes are like those in NPC (Kamranvar *et al*, 2007; Lacoste *et al*, 2010).

NPC and Burkitt’s lymphoma are thus two model EBV-related cancers. Both have increased chromosome structural changes with centromere involvement (Cuceu *et al*., 2018(Cu-ceu *et al*, 2018). Even a single break during these structural changes can cause breakage, fusion, and bridging. These three events then repeat during every cell division to destabilize the entire human genome. The chromosomes shatter, producing many more breaks (chromothripsis) and further complex rearrangements. Chromosome fragmentation is inherent to breakage, fusion, and bridge cycles (Umbreit *et al*, 2020) and occurs in all cancers much more frequently than previously thought (Cortes-Ciriano *et al*, 2020).

Robust, coordinated whole genome sequencing studies have delved deeply into the characteristics and origins of cancer genomes (Aleksandrov *et al*, 2020; Batra *et al*, 2021; Bruce *et al*, 2015; Cortes-Ciriano *et al*, 2020; Li *et al*, 2020; Meng *et al*, 2018; Nik-Zainal *et al*, 2016; Nones *et al*, 2019; Ou, 2020; Stephens *et al*, 2012; Voronina *et al*, 2020). These landmark studies provide vital starting points for the present work to compare breakpoints in breast cancers to breakpoints in cancers with proven relationships to herpes viruses.

## Results

### Viral homologies around breakpoints in *BRCA*. - associated breast cancers cluster around breakpoints in NPC, a model cancer caused by EBV

Every chromosome in female breast cancers has breakpoints that cluster near those in nasopharyngeal cancer, a known EBV-mediated cancer. This conclusion comes from analyses of genomes from 139 hereditary or familial breast cancers and 70 nasopharyngeal cancers (NPC). Fig. 1A shows the exact distances between NPC and breast cancer breakpoints that are within 200,000 base pairs of each other. These nearest neighbor breaks cluster around a few low valleys that periodically occur across the entire chromosome.

**Fig. 1A.**
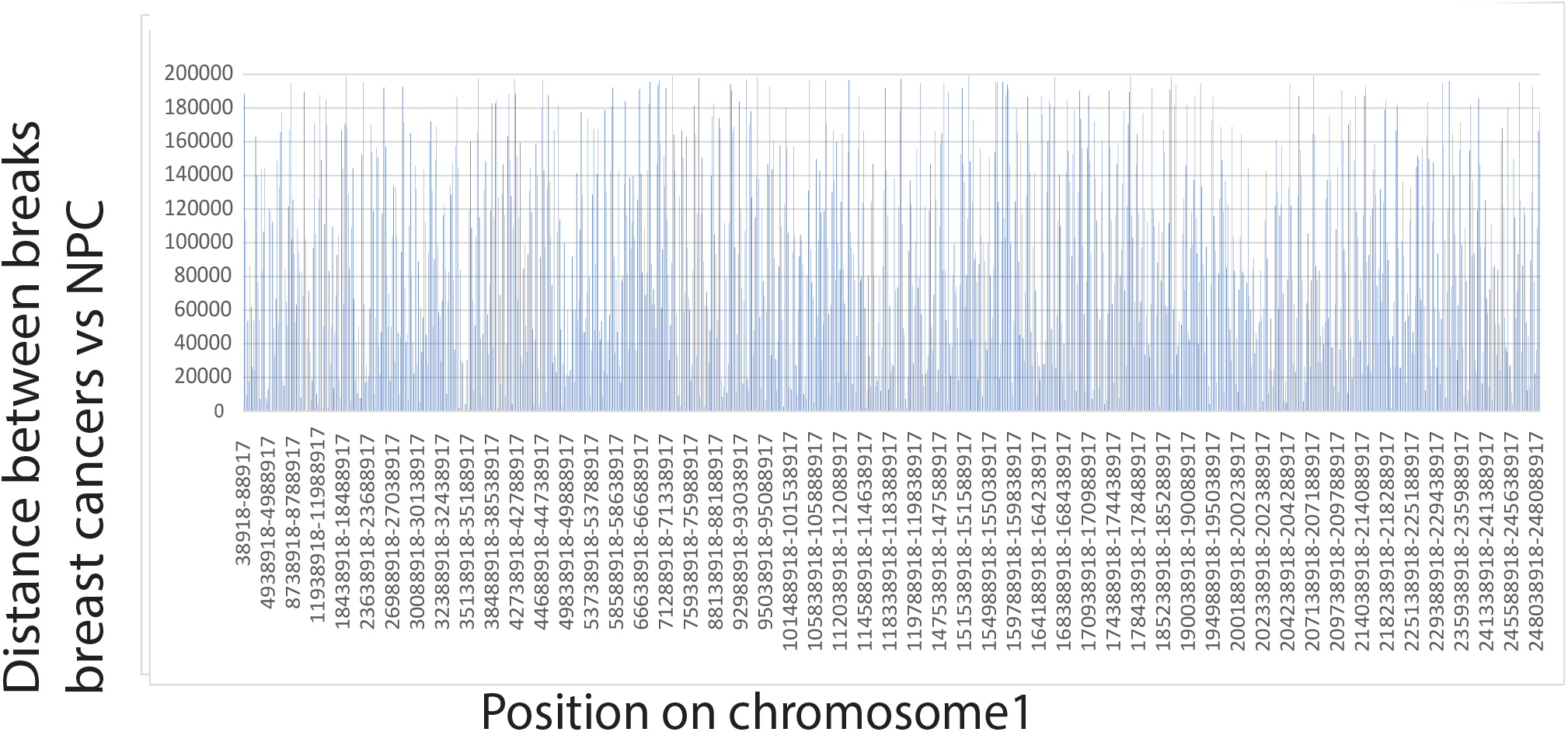
The exact distances between NPC and breast cancer breakpoints that are within 200,000 base pairs of each other.

Including the exact positions of all breakpoint pairs further than 200,000 base pairs apart results in a smeared graph that is difficult to interpret. However, cancers are collections of acquired mutations, indels, and longer chromosome copy number variations from different individual genomes. Moreover the data was collected over many years in different laboratories. Because of these complications, increments of 5000 base pairs were used to test for cyclic low valleys and to measure them in relation to all breakpoints (Fig. 1B). The 5000 base pair increment represents about 2e-5 error in relative positions for chromosomes 1 and 2. Some chromosomes shorter than chromosome 1 or chromosome 2 show the effect down to a 1000 base pair window. NPC vs breast cancers show a pronounced preference for breakpoints within 5000 base pairs but the exact percentages of breaks near each other is highly sensitive to the window size used. For 50 million base pairs on chromosome 22, 37% of the breaks occur within 175000 base pairs of each other and 5% of the breaks occur within 5000 bases. Many more breakpoints are well within the limits of the length of EBV variants (~175,000 base pairs). Table S1 presents all the calculations. Comparison of the plot for Chr2 in the top row vs the lower right shows this effect. The risk that any of these results is purely due to chance is less than 1 in 30,000, p<0.0001.

**Fig. 1B.**
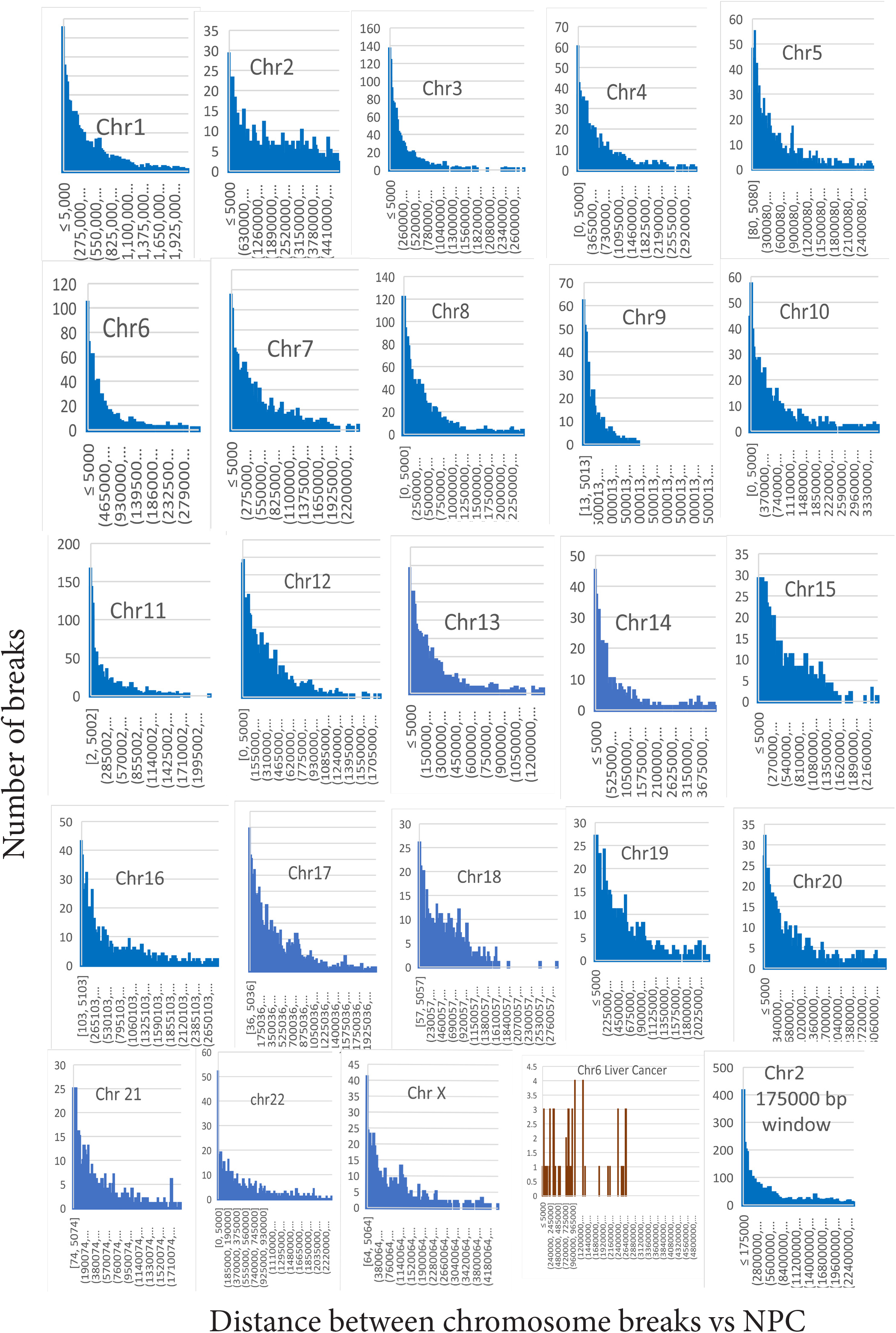
Breakpoints in 139 breast cancers that have a likely genetic contribution (BRCA mutation, familial, or early onset) cluster around the breakpoints found in 70 NPC’s. On all chromosomes, breakpoints in female breast cancers are most frequently within 5000 base pairs of breakpoints in NPC. As a control, an unrelated set of liver cancer data does not show the same relationship to NPC. On the X axis, each bar on the graph is separated by 5,000 base pairs but the drawing at the lower right shows how the selection of a larger bin size (the approximate length of EBV) affects the distributions of breakpoints.

A set of liver cancer breakpoints (Zheng *et al*, 2021) bear minimal resemblance to either breast or NPC breakpoints (Fig 1B, lower right). No breaks in 114 liver cancers on chromosome 1 were within 5,000 base pairs of breaks in any NPC, but one break in 61 liver cancers fit this window for chromosome 6. The chance that breakpoints on chromosomes 1, 2, 6, and 8 were more likely not within 5000 base pairs in liver vs NPC was 4.4 [CI=1.9-10] and were different from 1 (p=0.0004).

### Breakpoints in Burkitt’s lymphoma are near EBV variant sequences

Endemic Burkitt’s lymphoma provides another model cancer caused by EBV infection and tests whether results from NPC are general. Burkitt’s lymphomas have gene rearrange-ments at the Myc gene on chromosome 8, with variations in different patients (Busch *et al*, 2007; Walker *et al*, 2014). Fig. 2 focuses Chromosome 8 human-virus homology data on a 1.1 million base pair region around the Myc locus. There is a strong similarity in breakpoints in NPC vs breakpoints Burkitt lymphoma. Of 14 positions in NPC compared to Burkitt lymphoma breaks, 4 were identical and 3 additional breaks were within 2215 base pairs. Breast cancer comparisons also showed breakpoint positions similar to those in NPC in the same region with 12 values matching 32 Burkitt’s lymphoma breaks within 5000 base pairs (Fig. 2). The chance (relative risk) that these statistics differed from liver cancer was 14.9 (p=.0012).

**Fig. 2.**
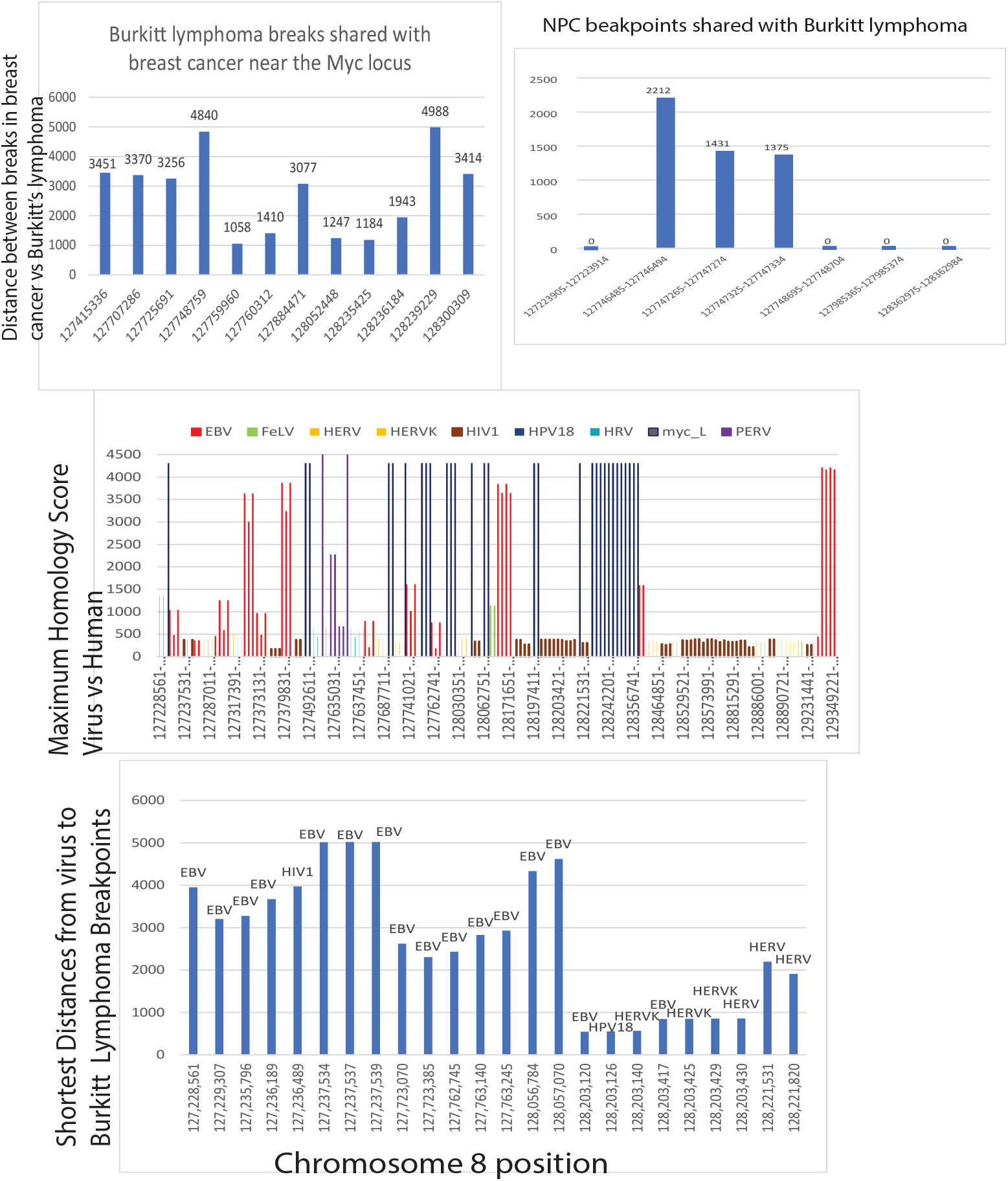
A 1.1 million base pair region around the Myc locus has a strong similarity in positions of breakpoints in NPC vs Burkitt lymphoma. Breast cancers have breakpoint positions similar to those in NPC in the same region with 12 breast cancer breaks matching 32 Burkitt’s lymphoma breaks within 5000 base pairs. Of 14 positions in NPC compared to Burkitt lymphoma breaks, 4 were identical and 3 additional breaks were within 2215 base pairs. The chance (relative risk) that these statistics differed from liver cancer was 14.9 (p=.0012). In the bottom panel, multiple DNA segments that strongly resemble EBV surround MYC locus breakpoints in Burkitt lymphoma. EBV predominates among the viruses closest to the breakpoints, but endogenous retrovirus and HIV1 sequences are also nearby. HPV18 near one break-point is unusual

Multiple DNA segments that strongly resemble EBV surround MYC locus breakpoints in Burkitt lymphoma. EBV predominates among the viruses closest to the breakpoints, but its presence does not absolutely predict a breakpoint. Human endogenous retrovirus (HERV) and human immunodeficiency virus (HIV1) sequences are also near some breaks. Human papilloma virus 18 (HPV18) near one breakpoint is unusual (Fig. 2).

### Breakpoints occur near human sequences that resemble herpes viruses

Virus-human homology comparisons were conducted around thousands of human BRCA1, BRCA2, and familial breast cancer-associated breakpoints using public data (Nik-Zainal *et al*, 2016; Nones *et al*, 2019). Long stretches of EBV variant DNA from two human gamma-herpesvirus 4 variants, HKNPC60 and HKHD40 are virtually identical to human breast cancer DNA around many inter-chromosomal breakpoints. Maximum homology scores for human DNA vs. herpes viral DNA over 4000 are common, representing 97% identity for nearly 2500 base pairs, with E “expect” values (essentially p-values) equal to 0. Breakpoints in hereditary breast cancers cluster around human sequences that resemble virus variants, especially EBV-like viruses. For example, Fig. 3 shows all viral homologies on the 145,138,636 base pairs of chromosome 8. Over 44,000 regions have significant homology to viruses with homology scores above 250, but only a few different viruses match these regions. This includes about 11,000 matches to EBV variants and 6000 matches to endogenous retroviruses. These homologies are not uniformly distributed among human chromosomes. Chromosome 17 (Fig.3, lower panel) reverses the relative numbers of human positions that match EBV (1462) vs endogenous retroviruses (10,445).

**Fig. 3.**
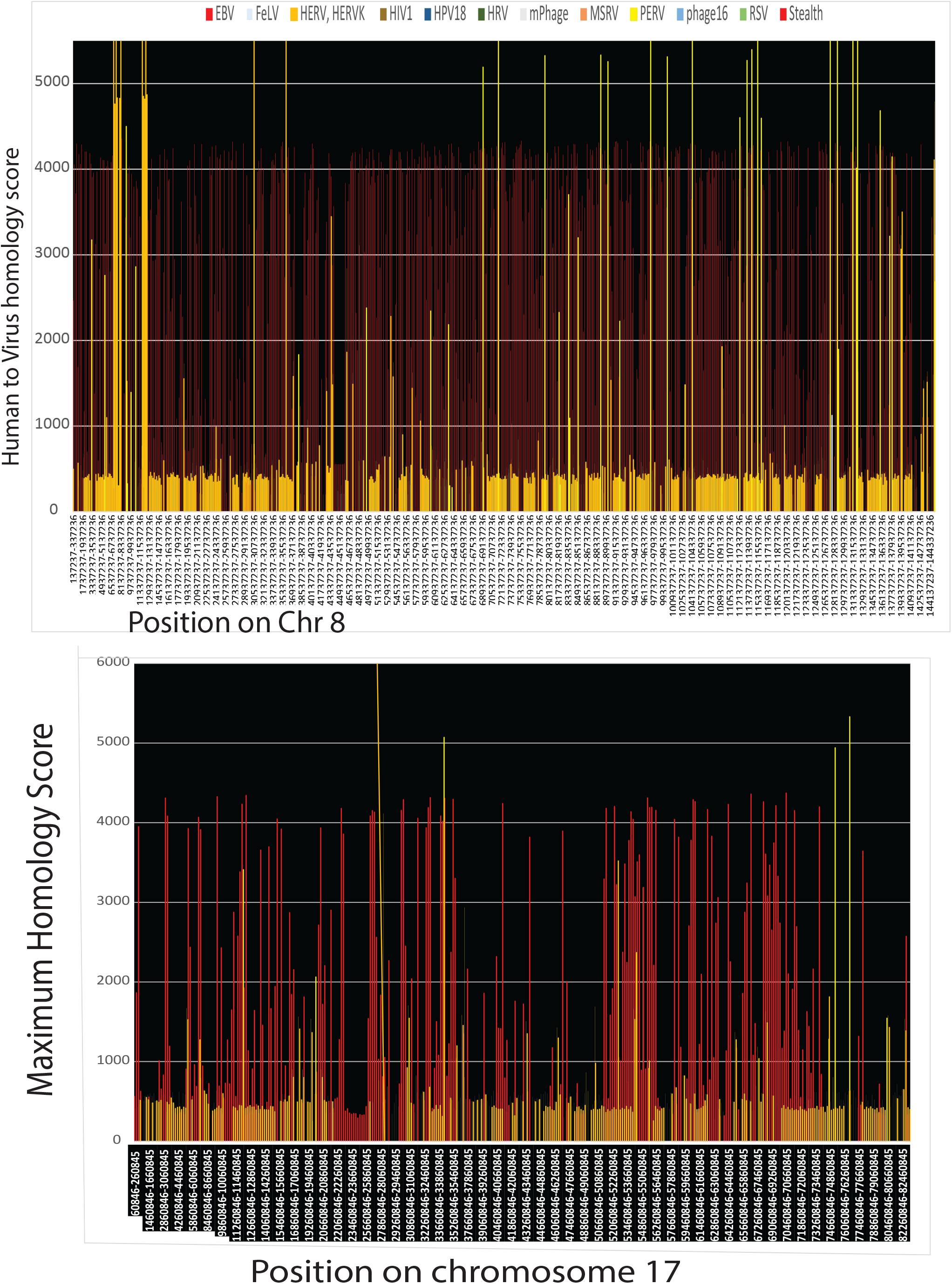
All viral homologies on the full lengths of chromosome 8 (top)and chromosome 17 (bot-tom). The two chromosomes have obviously different distributions of viral sequences. Chromo-some 8 is much more densely populated by virus-like sequences.

The positions of breaks in hereditary breast cancers aggregate near the position of human sequences homologous to viruses. On chromosome 8, 6823 viral sequences are within 5000 base pairs of a hereditary breast cancer breakpoint. On chromosome 17 there are only 1730 viral matching sequences within 5000 base pairs of a breast cancer breakpoint.

### Chromosome shattering clusters around sequences with viral homologies

The breakage-fusion-bridge cycle is a catastrophe related to telomeric dysfunction or double strand breaks. These events promote end-to-end fusions. During tumor generation, the cycle causes clustering of chromosome breakpoints, chromothripsis (Leibowitz *et al*, 2015; Umbreit *et al*, 2020), and aberrant repair with kataegis.

Fig 4 shows regions that undergo chromosome shattering with high confidence (Cortes-Ciriano *et al*, 2020) on the entire length of chromosome 6. There is a pronounced association of shattered breaks with viral sequences, especially for one patient. The two variables are strongly correlated by simple linear regression analysis (r2=0.93). To test this result, hierarchical cluster analysis was performed. High confidence chromothripsis positions or nearby matching viral sequences found either variable to be well described by two clusters. K-means analysis found the two chromothripsis cluster centers at 52,295,211 and 133,851,191 base pairs. Clusters of nearby viral matching sequences were almost the same (52,296,517 and 133,851,640). Very strong homology to EBV variants exists at both genome cluster centers. (Homology scores 3800-4000).

**Fig. 4.**
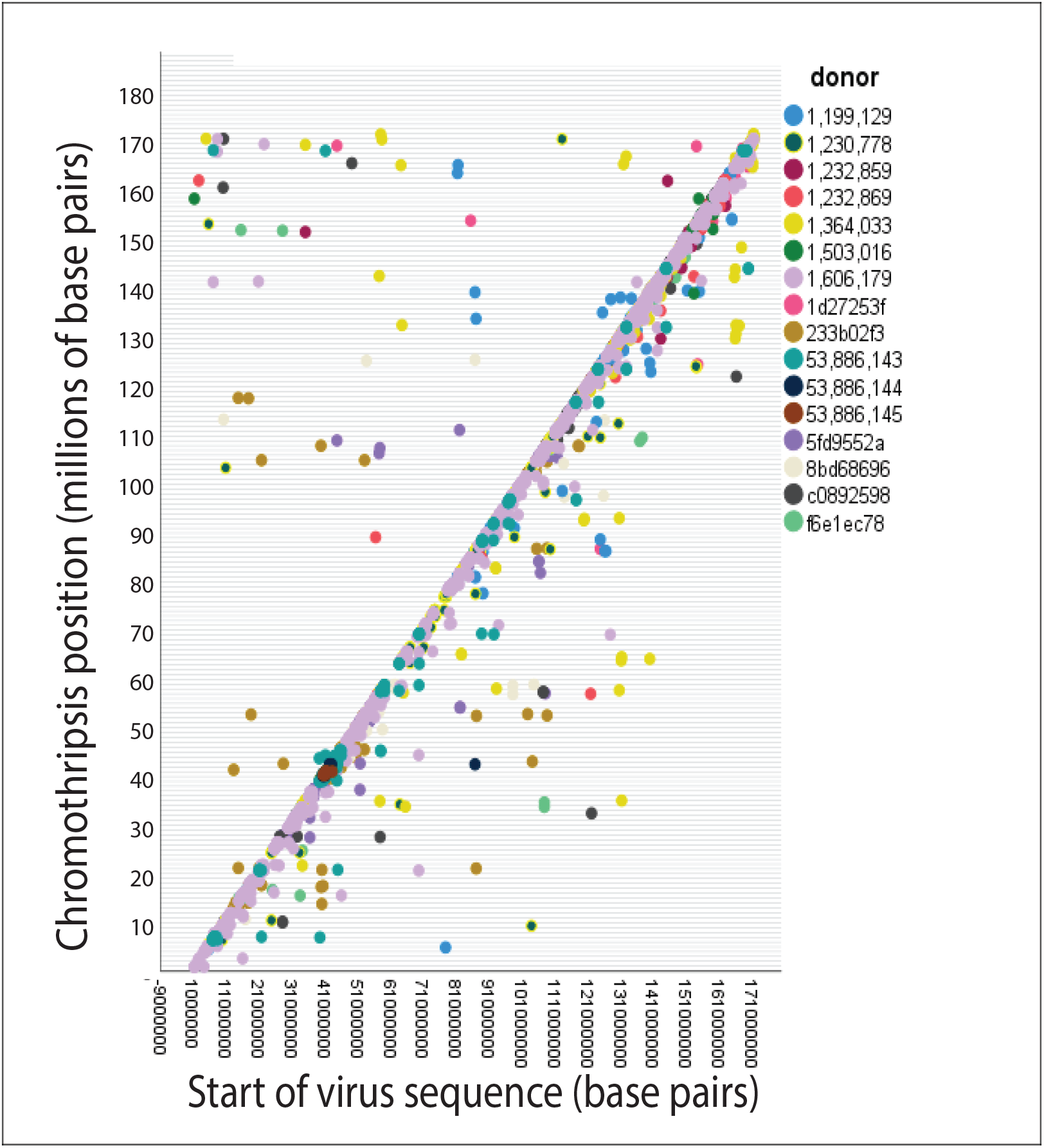
High-confidence positions where chromosome 6 shatters in 16 breast cancer genomes (Cortes-Ciriano *et al*, 2020) are plotted against start points of viral sequence homologies. The graph includes the entire length of chromosome 6 and implies a relationship between chromothripsis break-points and virus like sequences.

Conversely, 7918 positions of virus homology were grouped into 1000 clusters and the centers of the viral clusters were compared to chromothripsis breakpoints. Overall about 24% of the virus like cluster centers matched a chromothripsis breakpoint within 25,000 base pairs. Eleven virus like cluster centers matched chromothripsis breakpoints within 86-1000 base pairs. The eleven virus clusters contained 69 regions of virus homology with 57 matches to EBV variants.

Genome coordinates were graphically estimated for chromosome 6 fragments produced by high confidence chromothripsis events. Those fragments with copy numbers >=3 gave 1090 genome coordinates. Tests for normality showed these coordinates were unlikely to be a normal distribution (P<0.0001).

### Presumptive evidence of past EBV infection

Connections between EBV sequences and a damaged immune system provide presumptive evidence of past infection. Testing EBV sequences against all human sequences by BLAST analysis found about 65,000 areas of strong homology throughout the human genome (E<<e-10). Unlike retroviruses, EBV and its variants do not have integrase enzymes, so EBV/HHV4 should not integrate into the human genome. The presence of so many EBV sequences may be due to a human version of the bacterial CRISPR system.

Chromosome 6p21.3 contains MHC class 1 and class II genes, including HLA antigens HLA-A, HLA-B and HLA-C. Variants in these HLA antigens are strong risk factors for NPC infections (Tsao *et al*, 2017), because they metabolize antigens and present pieces of virus (viral antigens) to the immune system. Thirteen breast cancers in the COSMIC database have a deletion near the HLA region on chromosome 6 and about 23% of breast cancers have mutations directly affecting HLA class 1 or class 2 genes. Many more breast cancers have indirect connections with multiple damaged genes that interact with HLA antigens or are otherwise essential for immunity. Near the 5’ end of 6p21.3, these genes include MTR1, MOCS1, FANC-E, MTCH1, and ETV7. 139 breast cancers with a likely hereditary contribution have 284 breakpoints at chromosome 6p21.3.

In general, the piRNA system loosely resembles bacterial CRISPR. The piRNA system inactivates virus derived transposons related to human endogenous retroviruses by methylating or cleaving them. The distribution of viral sequences in the HLA region of chromosome 6p21.3 was compared to the distribution of piRNAs and transposons. Polymorphisms in HLA-DMB antigen at 6p21.3 have already been linked to the herpes virus mediated cancer Kaposi’s sarcoma (Aissani *et al*, 2014). Fig. 5A shows a marked similarity in how remnants of exogenous and endogenous viruses are distributed at 6p21.3. Both have homology to the same human sequence and are sandwiched between piRNA sequences, sometimes right next to each other. A 10,000 base pair window enables viewing the entire area (Fig. 5A). At this resolution every viral sequence is sandwiched between two piRNA sequences and grossly resembles the bacterial CRISPR system.

**Fig. 5A.**
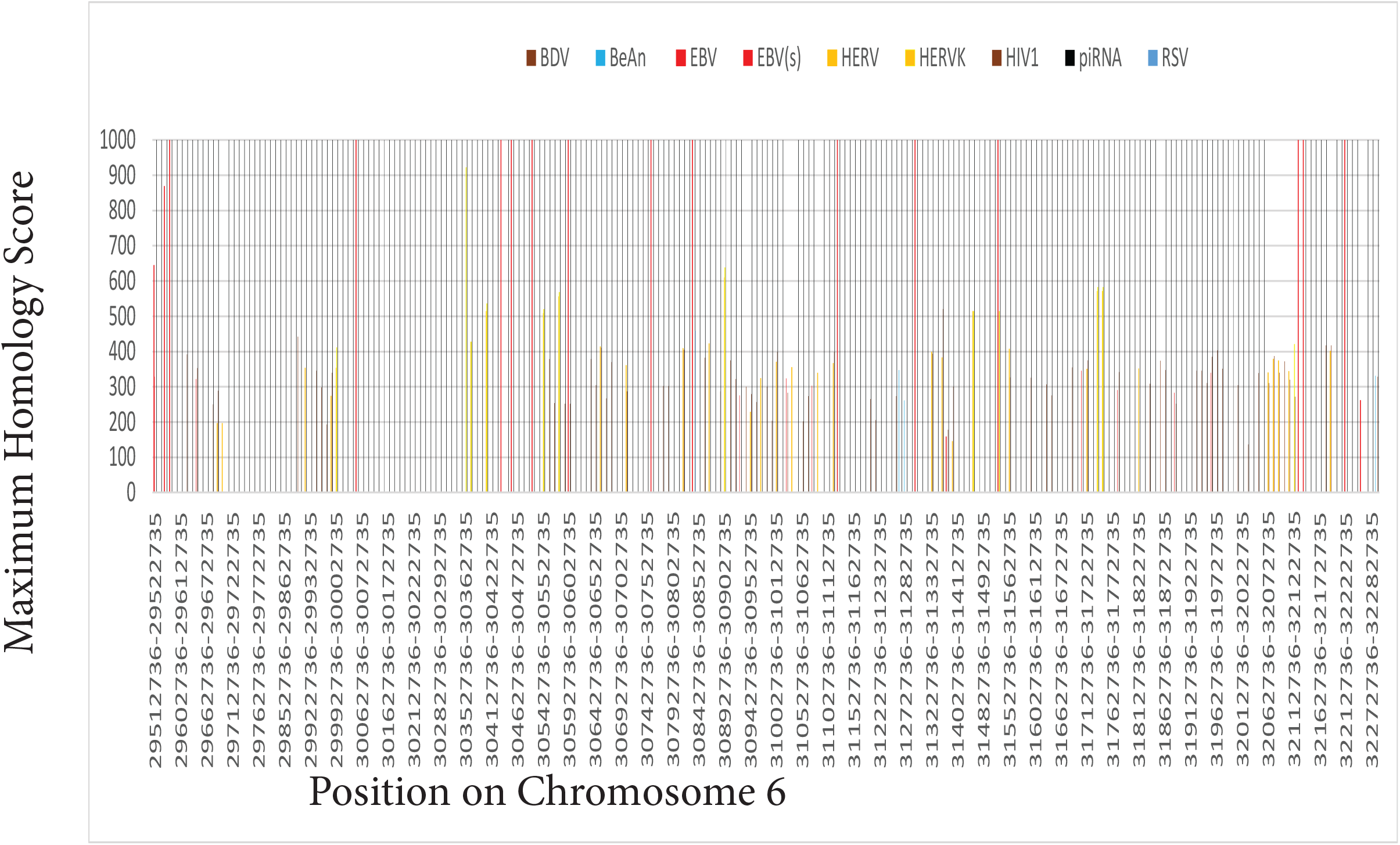
Remnants of exogenous and endogenous viruses distribute like each other over 2.77 million bases on chromosome 6p21.3. The data is best viewed at higher magnification. Both exogenous and endogenous viruses have homology to the same human sequence and are sandwiched between piRNA sequences, sometimes immediately adjacent.

HIV1 and HERV sequences are adjacent in numerous locations and sometimes EBV fragments are also present. This interspaced arrangement suggests piRNA defense mechanisms not only inactivate transposons related to endogenous viruses but also exogenous viruses, such as EBV variants. Long stretches of endogenous transposon-like DNA sequences are almost identical to exogenous viruses. A random sample demonstrates this homology for endogenous transposons (HERV) and exogenous viral sequences (HIV1, Stealth virus 1, BeAn 58058, HPV16, and Zika) (Fig. 5B). However, HERV sequences are more abundant than exogenous viruses, so there are regions where HERV transposons are the only ones present.

**Fig. 5B.**
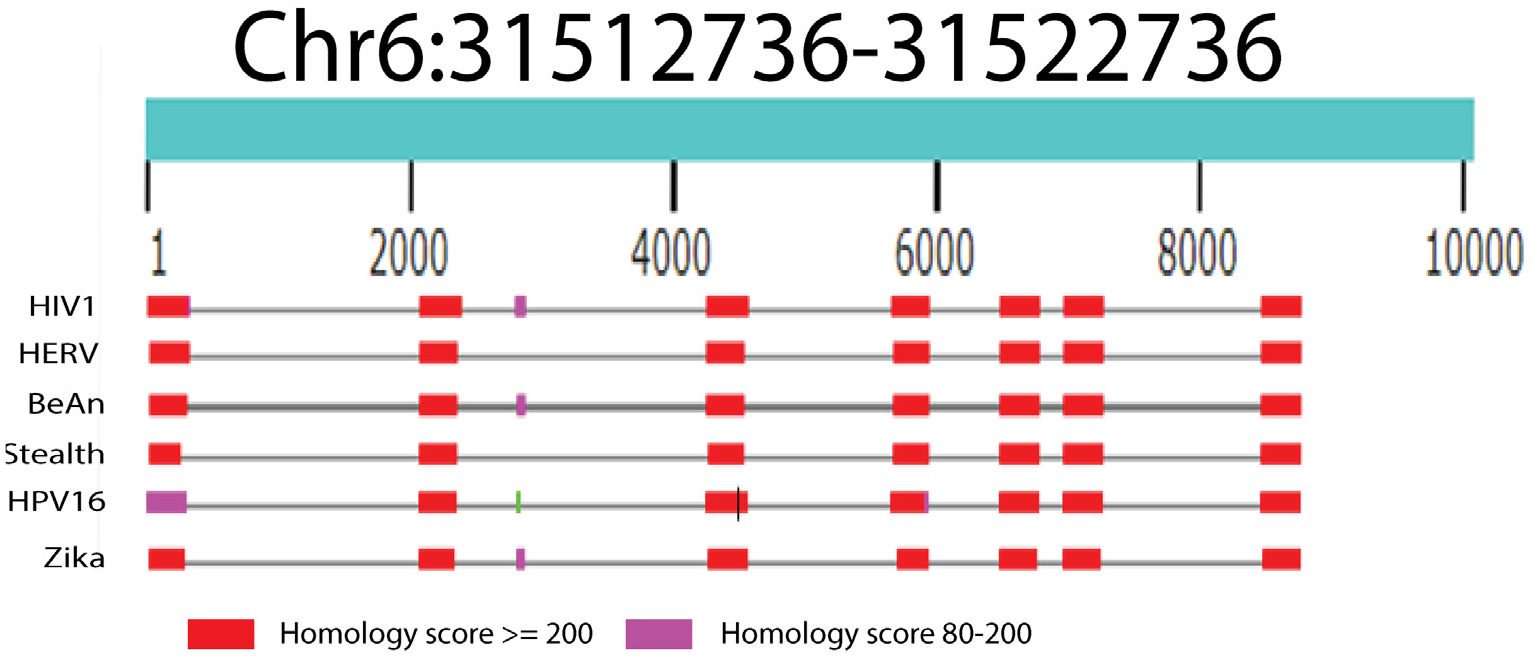
Transposons as endogenous retrovirus fragments are not easily distinguished from exogenous viruses because incorporated fragments strongly resemble each other.

Levels of various piRNAs differ by more than 1000-fold, but only the most abundant piRNAs are present in every cell. These abundant sequences drive the inactivation of foreign DNA. These rare piRNAs cannot function in every cell, but can potentially adapt to new genome invaders (Genzor *et al*, 2021). The human genome organizes piRNA sequences into clusters (Girard *et al*, 2006). Fig. 5C shows piRNA sequences clustered near the MHC region of chromosome 6 (at ~30-40 Megabases) with hundreds of piRNAs in close proximity.

**Fig. 5C.**
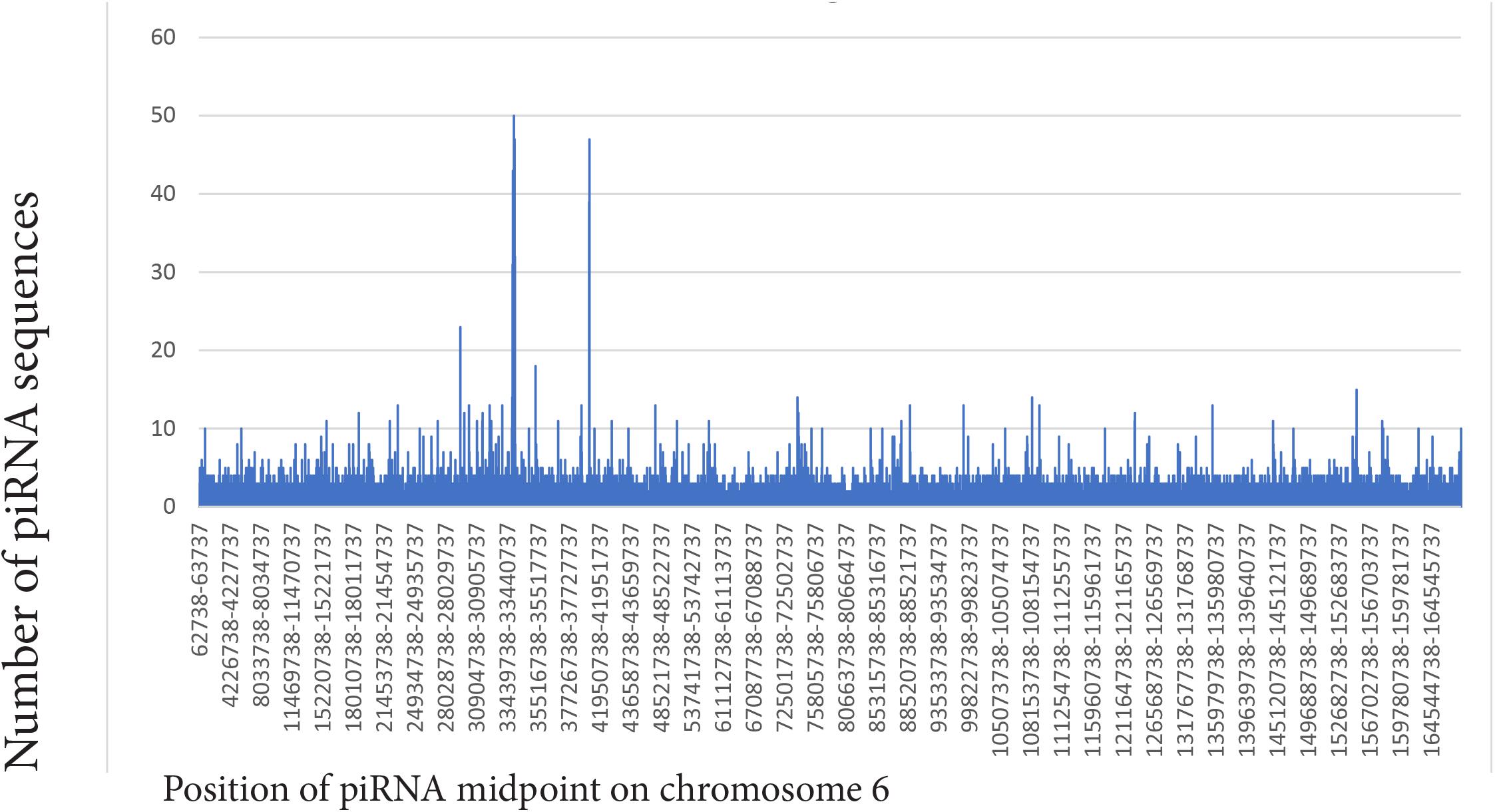
piRNA sequences are abundant near chromosome 6p21.3. Clusters containing hundreds of piRNAs are not present anywhere else on chromosome 6.

Inactivation of genes by hypermethylation at chromosome 6p21.3 was tested and found near the clustered piRNAs. While EBV related gastric cancer and NPC have distinct patterns of DNA hyper-methylation, some chromosome areas are hypermethylated in both cancers. Chromosome 6p21.3 is one of these areas and contains a potential EBV infection signature (Scott, 2017). This potential marker was examined in 1538 breast cancers using existing methylation data (Batra *et al*, 2021). Fig. 5D shows that this marker region in breast cancers has significant differences in promoter methylation vs. normal controls. Promoters for the genes in Fig. 5D inhibited by hypermethylation prevent tumors (TNFB) (Fernandes *et al*, 2016), modulate cancer progression (SCUBE3) (Wu *et al*, 2011) and respond to antigen-antibody complexes (C2).

**Fig. 5D.**
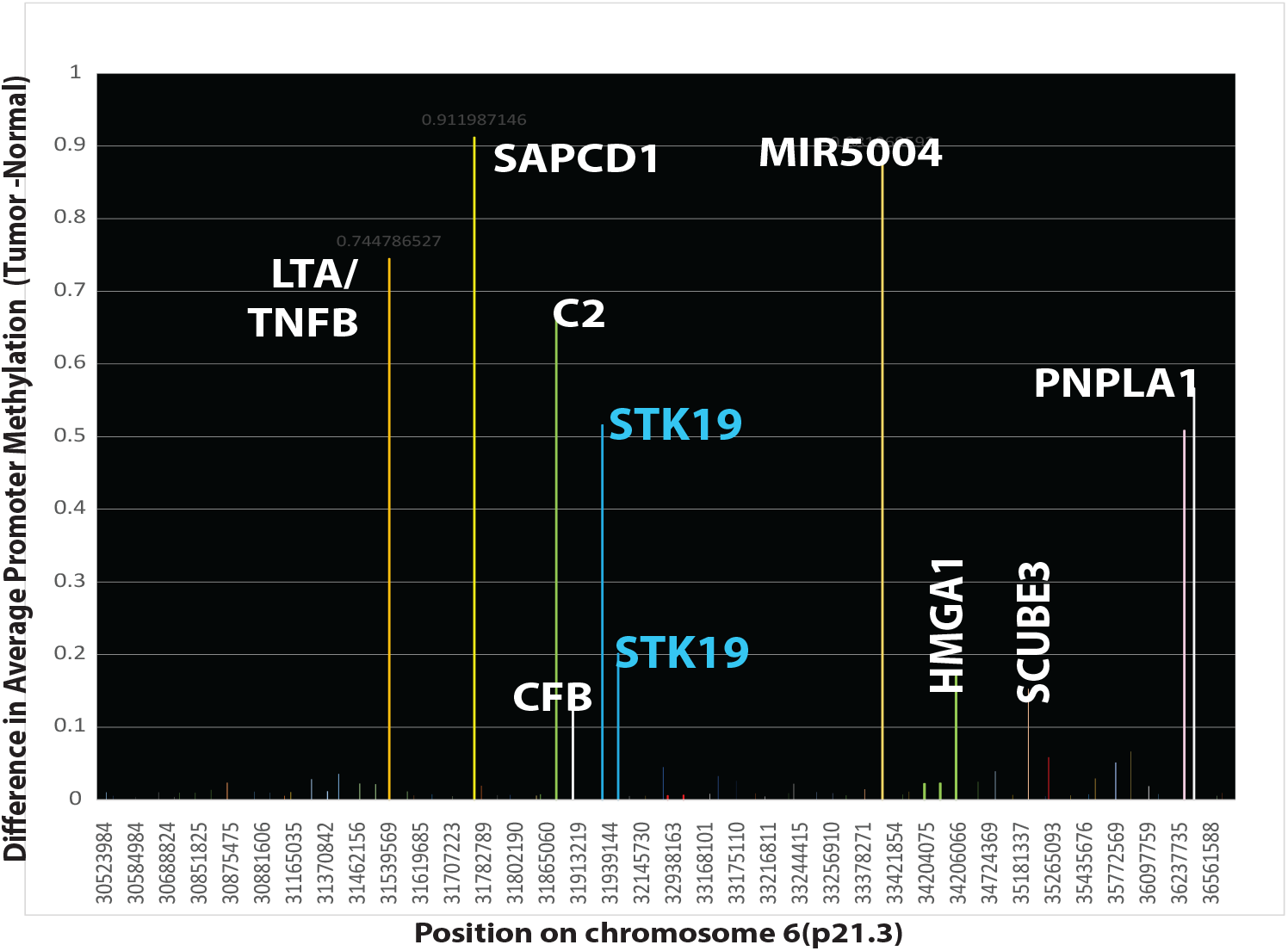
Chromosome 6p21.3 also contains a potential EBV infection signature (Scott, 2017). This marker region in breast cancers has significant differences in promoter methylation vs. normal controls.

Extending the comparisons of viral sequences and piRNAs down to a 50 base pair resolution reveals additional details (Fig. 5E). At this resolution, a few short EBV variant sequences fit within the borders of two piRNAs (darker red areas). Longer EBV variant sequences encompass one or two piRNAs, so that the piRNAs can derive from the dead virus.

**Fig. 5E.**
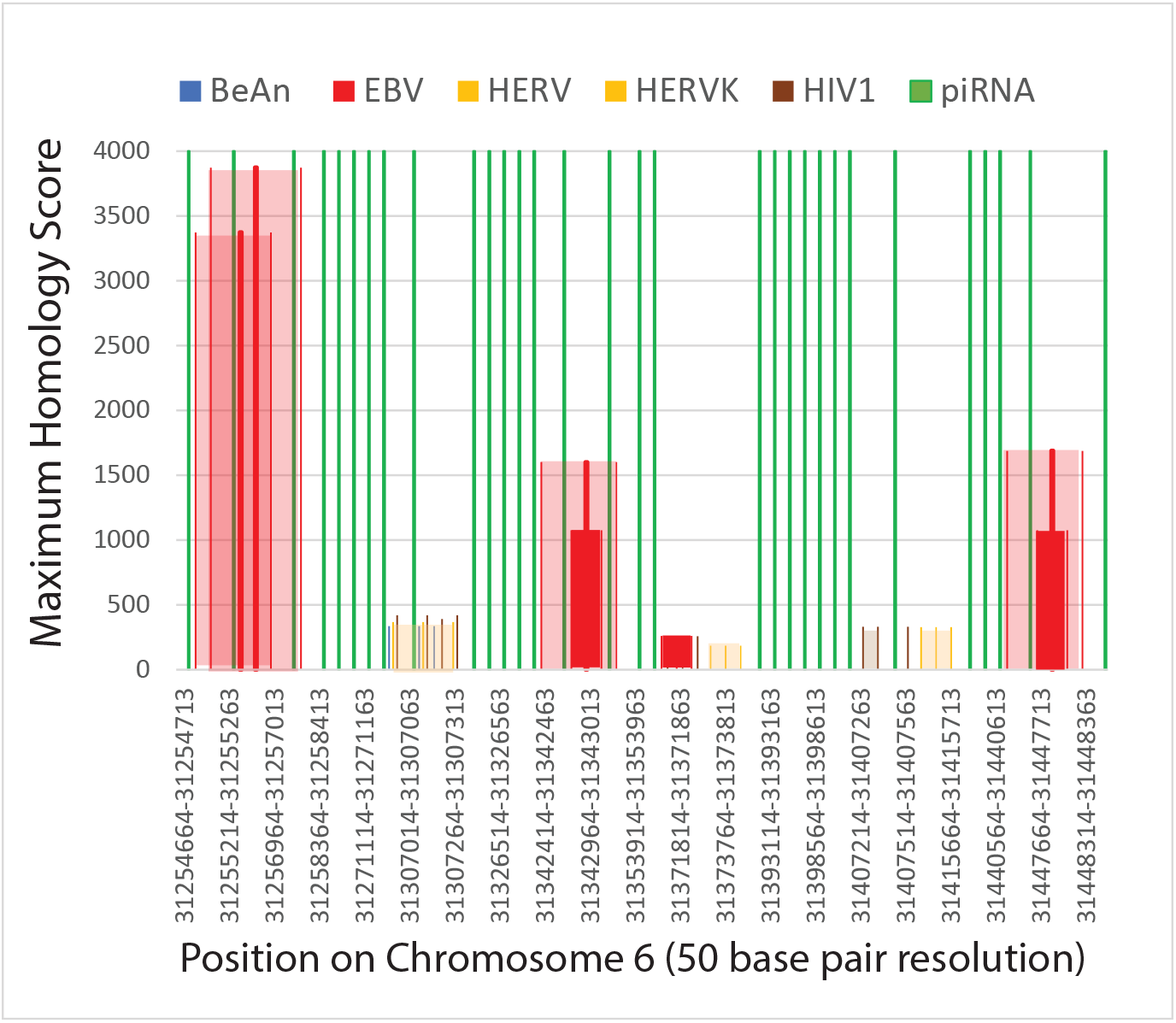
High resolution 50 base-pair comparison of piRNAs to regions of viral homology.

In contrast to piRNAs present throughout chromosome 6, other regions of the human genome apparently do not have piRNA sequences. For example, even though chromosome 17 also contains piRNA clusters (Girard *et al*, 2006), a virus rich region of chromosome 17 from about 50.00-56.47 million bases does not have piRNAs, so the viral sequences incorporated there are not subject to piRNA-type controls.

### An Identified EBV genome docking site is near breast cancer breakpoints

An independent study has isolated evidence of a proximity of breast cancer breaks to an EBV genome docking site on chromosome 11. One primary EBV genome docking area on chr11 near the FAM-D and FAM-B genes (Lu *et al*, 2010) is close to two breast cancer breakpoints (Fig. 6). On chromosomes 1 and 5, other breast cancer breaks are further from EBV docking sites but still well within the length of the EBV virus sequence.

**Fig. 6.**
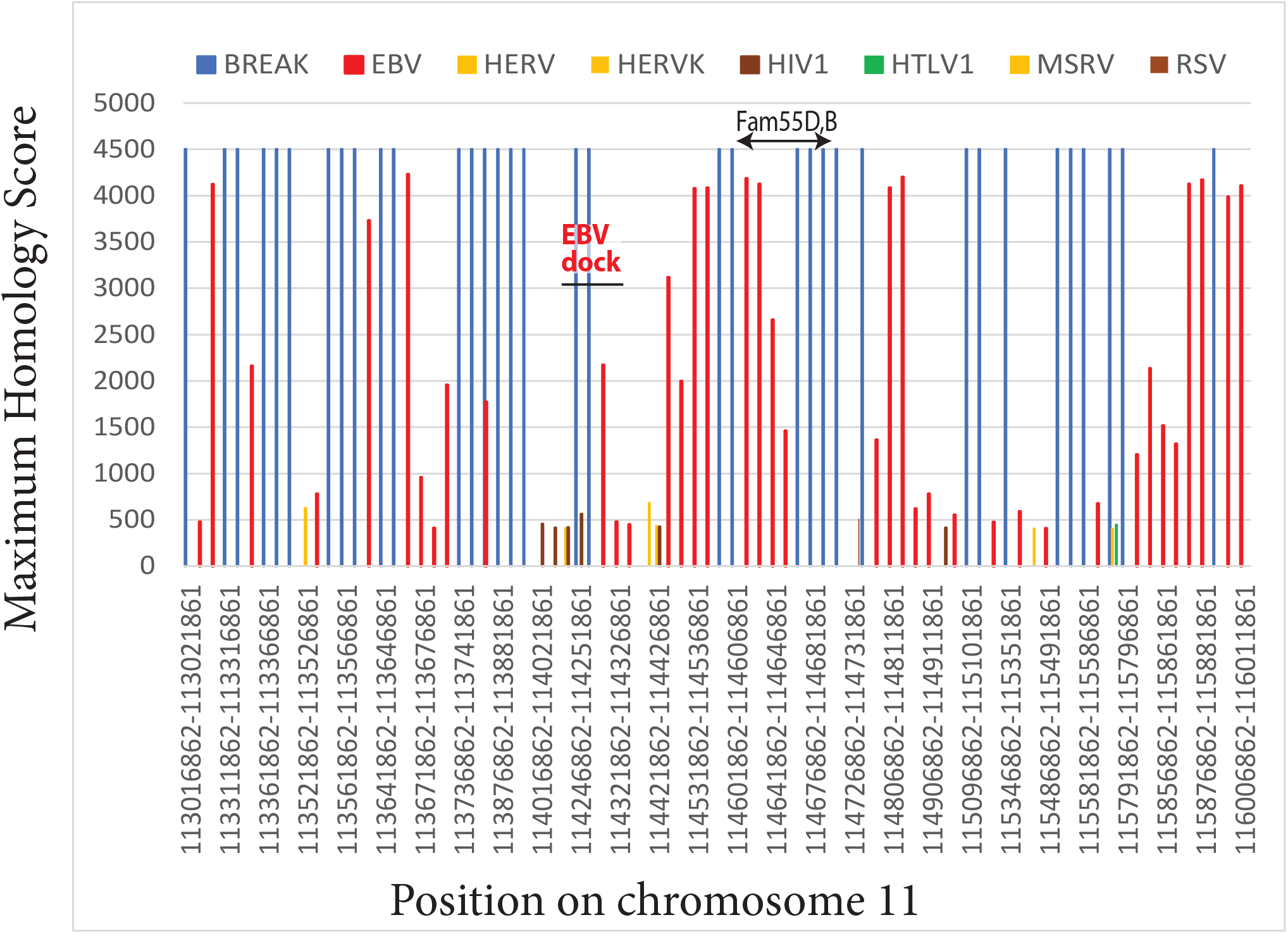
Maximum homology to human DNA for all viruses (y-axis) is plotted for known EBV genome anchor sites on chromosome 11 vs. breast cancer breakpoints and gene coordinates. A primary EBV docking site (Lu *et al*, 2010) overlaps two breast cancer breaks and human DNA that resembles EBV variants.

### Testing alternate explanations

#### BRCA1 or BRCA2 mutations by themselves are not sufficient to cause chromosome breaks

The possibility was tested that inherited mutations, such as those in BRCA1 or BRCA2 genes, are sufficient to cause chromosome breaks without contributions from herpes variants or other viruses. Seventy-four women over age 70 with normal BRCA genes and no other known inherited cancer-associated mutations (Nik-Zainal *et al*, 2016) were taken as a sporadic breast cancer group. Breakpoints in their cancer genomes were compared to breakpoints in breast cancers with a likely genetic component (e.g. a known BRCA mutation, elevated risk for an inherited gene mutation, or early onset). Although there are significant differences in breakpoint distributions, many breakpoints apparently cluster at similar chromosome locations (Fig. 7A).

**Fig. 7A.**
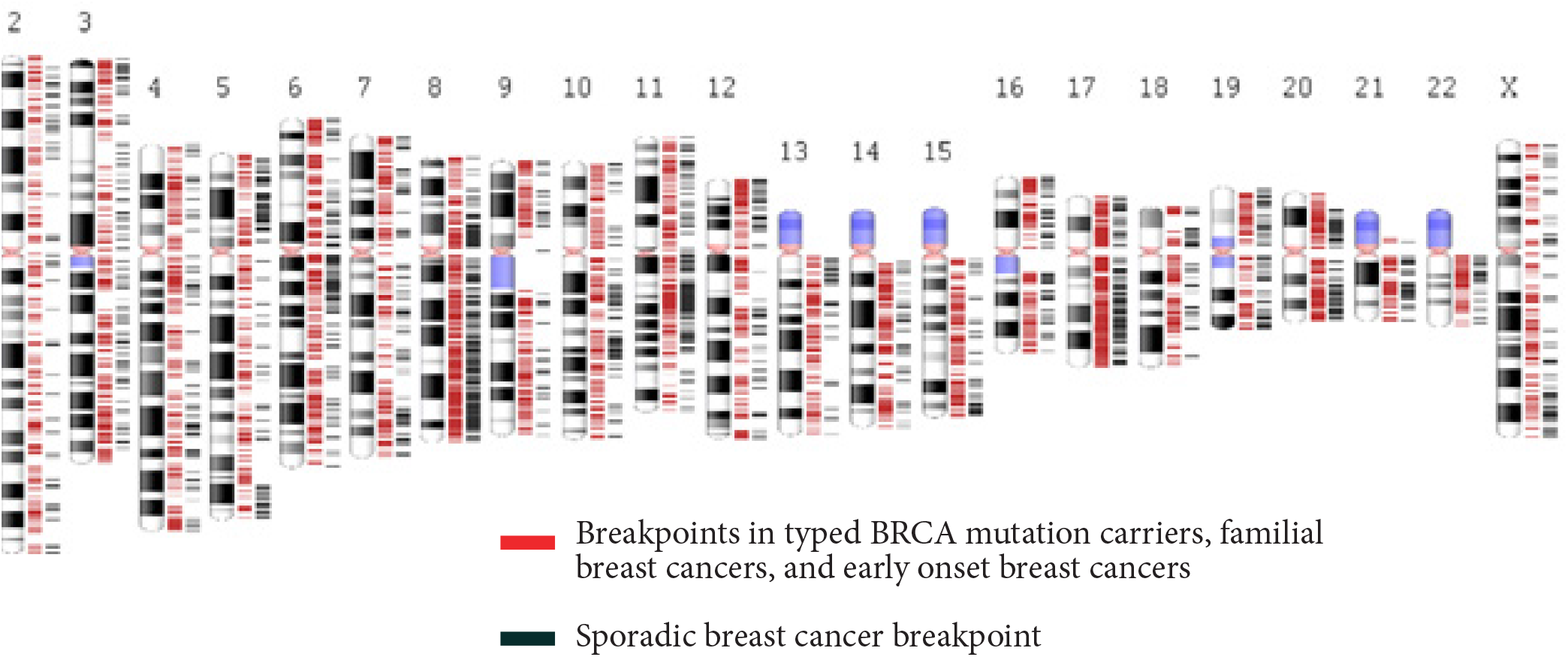
Inter-chromosome translocation break positions in 74 mutation-associated, familial, or early onset female breast cancers (red) vs. 74 likely sporadic female breast cancers (black).

To test this result further, NPC was again used as a model for EBV-mediated breakages. Breakpoint positions in the 74 sporadic breast cancers were compared to breakpoints in 70 nasopharyngeal cancers. Breakpoints within 5000 base pairs of a break in NPC are the largest single category on most chromosomes (Fig. 7B). Chromosome 11 has 9% of its breaks at a distance <=5000 base pairs from an NPC break and almost all its breaks (76%) are within the length of EBV (~175, 000 base pairs) from an NPC break. Chromosome 22 has 19% of its breaks within 5000 base pairs of an NPC breakpoint and 46% of its breakpoints within 175,000 base pairs. In contrast on other chromosomes there are significant numbers of breast cancer breakpoints more distant (>5000 base pairs) from NPC breakpoints (e. g. chromosomes 9,10, and 18). These analyses support an association between sporadic breast cancer breaks and breakpoints in NPC. This association occurs without BRCA1 and BRCA2 gene mutations.

**Fig. 7B.**
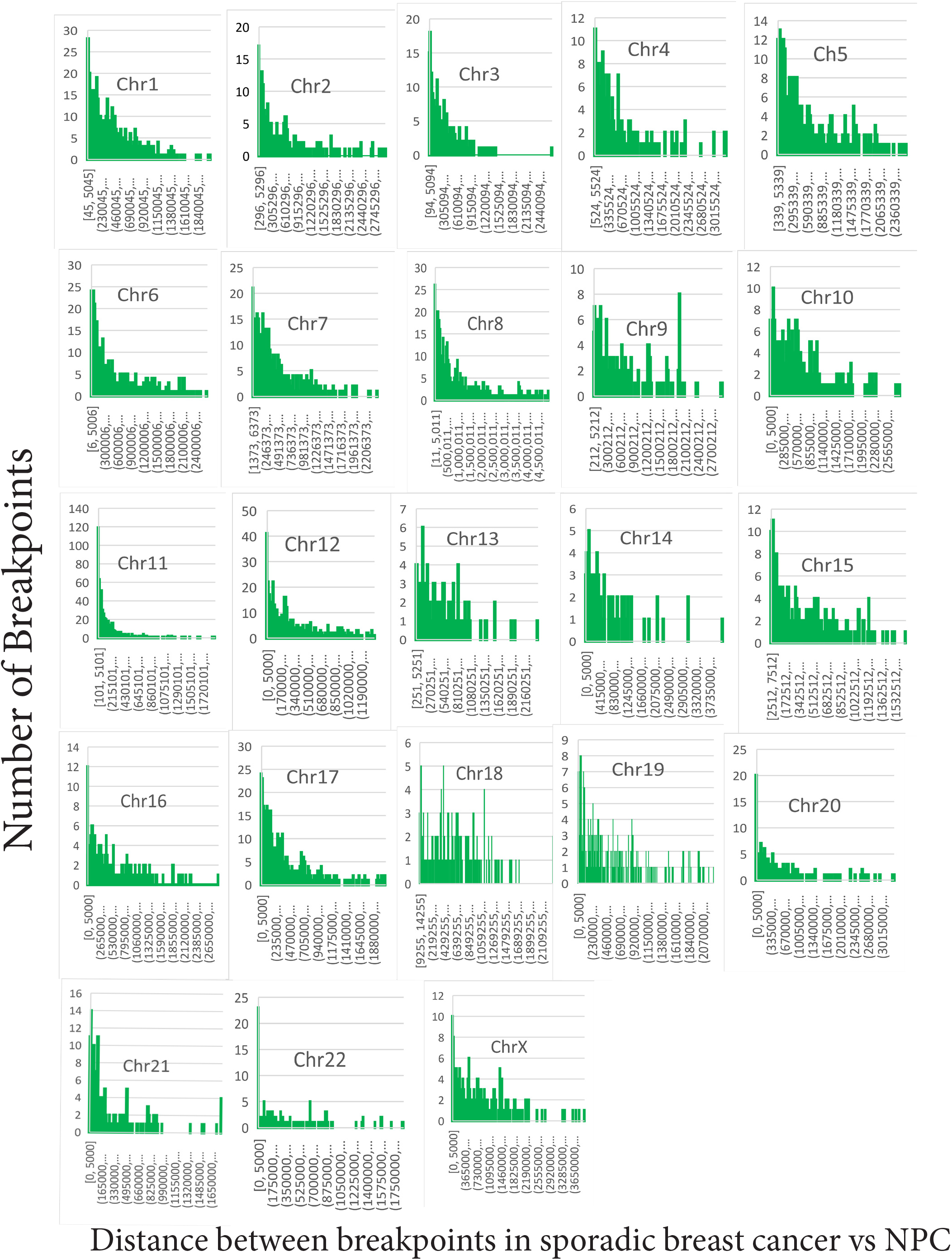
Breakpoints in 74 sporadic breast cancers cluster around the breakpoints found in 70 NPC’s. On all chromosomes, breakpoints in female breast cancers are most frequently within 5000 base pairs of breakpoints in NPC.

Promoter hypermethylation is a potential confounder because it provides an alternate method of inactivating BRCA1 and BRCA2 genes. BRCA1 and BRCA2 methylation is unusual in sporadic cancers. Average methylation scores in 1538 sporadic breast cancers (Batra *et al*, 2021) were 0.050 and 0.004 for BRCA1 and BRCA2 promoters, respectively. Triple-negative breast cancer may be another significant confounder because up to 58.6% of 237 patients had a marker predictive of BRCA1/BRCA2 deficiency (Staaf *et al*, 2019). However, triple-negative breast cancers com-prised only 9% of breast cancers in the cohort.

#### Human sequences do not explain virus matches

An alternate explanation for DNA sequences matching herpes viruses is that they are a repetitive sequence in the human genome unrelated to viruses. To test this explanation HKHD40 and HKNPC60 were compared to all herpes viruses. About 100 other gamma herpes viral variants strongly matched HKHD40 and HKNPC60 in regions with enough data to make comparisons possible. HKNPC60 is 99% identical to the EBV reference sequence at bases 1-7500 and 95% identical at bases 1,200,000-1,405,000. HKHD40 has 99% and 98% identity in the same regions. HKHD40 and HKNPC60 are typical of many other herpes-virus isolates, with some haplotypes conferring a high NPC risk (Xu *et al*, 2019).

#### Other viruses also contribute

The total numbers of EBV like sequences in human were estimated by comparing HKHD40 and HKNPC60 sequences to all human sequences. This produced nearly 65,000 matches. Other viruses implicated in breast cancer such as MMTV and HPV16 rarely matched human chromosome segments. Testing the retrovirus MMTV produced 12 significant matches to the human reference genome and the maximum bit score was 1164 (small by comparison to other viruses). MMTV also has a 22 bp match to a retrotransposon on chromosomes 2, 19, 18 and 17, but MMTV has significant homology with a retroviral gag-pol-pro precursor gene found in some human breast cancers (Liu *et al*, 2001). Retroviral sequences constitute 8% of the human genome. EBV can trans-activate endogenous retroviruses (Bruce *et al*, 2021; Mameli *et al*, 2012; Meier *et al*, 2021), In the present work, I did not address how retroviruses impact breast cancer risk, but they may make a significant contribution.

#### Breakpoint positions are not random events

The possibility was tested that breakpoint positions result from random, independent events. From probability theory, the sums of independent random variables tend toward a normal distribution (i.e., a bell-shaped curve) even if the original variables themselves do not fit a normal distribution. Then breast cancer breakpoints should follow a Gaussian distribution. Contrary to this prediction, statistical tests for normality find no Gaussian distribution for thousands of breakpoints in any breast- or nasopharyngeal cancer on any chromosome (p<0.0001). Instead, many breakpoint positions are approximately the same among different breast and NPC cancers.

#### Fragile site breaks do not account for breast cancer breakpoints

Lu et al. found 4785 EBNA1 binding sites with over 50% overlapping potential fragile sites as a repetitive sequence element (Lu *et al*, 2010). Based on the fragile site database, chromosome 1 contains 658 fragile site genes, the most of any chromosome (Kumar *et al*, 2019). Although some fragile sites align with breast cancer breaks on chromosome 1, large numbers of breaks on chromosomes 4, 12 and most other chromosomes are inconsistent with the fragile site database. In the PCAWG dataset, there are no fragile sites listed for chromosomes 8, 9, 11-15,17-19, 21, and 22 (Li *et al*, 2020). There are many more breakpoints than fragile sites on all the chromosomes. Breast cancer breaks do not consistently occur near common fragile sites. Some hereditary breast cancer breakpoints were tested for repeats likely to generate fragile site breaks because the repeats are prone to replication errors. This test did not find such sequences.

## Discussion

The present work shows human sequences related to tumor virus variants cluster around chromosome breaks in breast cancers. Virus like human sequences surround high-confidence chromothripsis breaks, which are an inherent feature of anaphase bridges. Multiple independent lines of evidence support this view and are summarized below.

1. There is a clustering of breast cancer breakpoints near breakpoints in NPC on all chromosomes in females, regardless of whether the breast cancers have a genetic component. NPC is a model for an epithelial cell cancer associated with EBV.
2. Breakpoints in Burkitt’s lymphoma, another model EBV cancer are also related to those in breast cancer.
3. Every human chromosome in female breast cancers has breakpoints near EBV-like human DNA. EBV can also activate endogenous retroviral sequences, found near some breakpoints.
4. Chromosome fragmentation is an integral part of breakage-fusion-bridge cycles. On chromosome 6, boundaries where chromosome fragmentation has occurred match positions of viral homology. Model EBV infected B-cells show chromosomes with two centromeres and other evidence of breakage fusion bridge cycles.
5. Endogenous retroviral transposons are inactivated by piRNAs in a system grossly resembling bacterial CRISPR. Relative to piRNAs, exogenous virus fragments are arranged just like endogenous retroviral transposons and some are highly homologous to them. This occurs in the MHC class I and class II region of chromosome 6.
  A nearby potential EBV methylation signature shared with known EBV-related cancers is far more abundant in 1538 breast cancers than in normal controls. Methylation is one method used by piRNA mechanisms to inactivate transposons.
6. Two areas where EBV docks to the human genome are near breakpoints
7. The association of breast cancer breaks and EBV tumor variants does not depend on the presence of BRCA1 or BRCA2 gene mutations. However, breast cancer breaks are more frequent in breast cancers that have a genetic component than in cancer that do not.
8. The association of EBV variant sequences with chromosome breakpoints does not require the continuing presence of active viruses. One breakpoint in a single cell can generate further breaks during cell division and destabilize the entire human genome. Clusters of mutations also occur after improper break repair.
9. Fragile site sequences are not sufficient to account for breast cancer breaks
10. Tumor variants HKHD40 and HKNPC60 are herpesviruses closely related to known tumor virus populations KHSV and EBV.

Some of the most recent cancer drug therapy focuses on identifying and targeting cancer driver mutations. The drugs are initially effective, sometimes for long periods but then the drugs quit working. The graphical abstract summarizes a proposed model showing breakage-fusion-bridge cycles as an occult, underlying process that continues even if cells with the initial driver mutations are removed. Cancer treatment generates new clones that do not exist in the original population (Aissa *et al*, 2021). The cancer phenome includes an inherent process that produces new cancer driver mutations and new cancer phenotypes (graphical abstract).

According to the model (Graphical Abstract), mutations and chromosome breaks affect immunity and damage control of EBV infection. The mutations affect processes such as cytokine production, autophagy, etc. One damaged gene can cripple an entire immune function, which relies on many genes dispersed throughout the genome. Each breast cancer genome can have different mutations affecting immunity, with the same gene only occasionally damaged in different cancers (Friedenson, 2013; Friedenson, 2015).

The piwi-piRNA pathway adds to the immune system another layer of protection for the genome. Endogenous and exogenous virus remnants are positioned among piRNAs on the MHC region of Chr6, near a methylation signature for EBV infection. The pathway silences transposons and retrotransposons such as those from endogenous viruses, reducing double-strand break formation in both germ and somatic cells (Mani and Juliano, 2013; Toth *et al*, 2016). The arrangement near HLA antigen genes facilitates communication between a system for DNA protection, innate, and adaptive immunity.

The piwi-piRNA pathway induces heterochromatin formation at centromeres, thus affecting how centromeres behave in breakage fusion cycles (Thomson and Lin, 2009). Centromere amplifications are common in breast cancer, accompanying HER2 amplification (Davies and Voutsadakis, 2020). Suppose a chromosome fragment includes a centromere but is deficient in telomeres, so a second centromere-containing fragment can link to it. The product now has two centromeres. During anaphase in cell division, the two centromeres try to separate, form a bridge, and the chromosome breaks again (McClintock, 1941). The bridge region does not replicate normally during mitosis, so chromothripsis and kataegis become characteristic of the process (Umbreit *et al*, 2020). This scenario occurs in EBV-positive Burkitt lymphomas (Kamranvar *et al*, 2007; Lacoste *et al*, 2010) and produces visible anaphase bridges in precursor forms of breast cancer (Chin *et al*, 2004). Chromosome shattering accompanying these bridges occurs in at least half of breast cancers (Cortes-Ciriano *et al*, 2020). Traditional chromothripsis involves alternate copy-number states for chromosome regions, but additional events involve multiple chromosomes and structural alterations. These alterations arise from non-homologous end joining, replication difficulty, and insertion of templated fragments. Chromothripsis can amplify oncogenes and inactivate DNA repair genes (Cortes-Ciriano *et al*, 2020). Positions of chromothripsis correlate well with positions of viral sequence homology.

Translocations can generate a burst of localized somatic mutations through the actions of APOBEC (“kataegis”) (Nik-Zainal and Morganella, 2017). APOBEC is typically another response to inactivate viral infections. Effects related to APOBEC3 probably occur in EBV-induced carcinogenesis (Bobrovnitchaia *et al*, 2020; Law *et al*, 2020). The breakage fusion bridge process repeats during cell division, quickly increasing the number of abnormal chromosome products and driver mutations. When telomeres are removed or become so short that they cannot protect chromosome ends, APOBEC3B cytosine deamination (and perhaps TREX1 nucleolytic processing) cause chromothripsis and kataegis (Maciejowski *et al*, 2020; Umbreit *et al*, 2020). It is difficult to imagine how a spurious EBV-like chromosome would not contribute to these events.

Removing malignant cells by targeting a driver mutation stops cancer, but this rarely involves 100% of cancer cells. As the remaining cells divide, the underlying process continues to produce more driver mutations and new phenotypes, so the cancer comes back. Once the process starts, it remains active without requiring large numbers of EBV particles. Chromothripsis, an integral part of this circuit has been linked to poor survival in cancers of the central nervous system, colorectal cancer, and acute myeloid leukemia (Voronina *et al*, 2020). Driver genes generated by chromoplexy (Shen, 2013) are also consistent with this model. The model does not rule out contributions from subpopulations of breast cancer cells that have differential sensitivity to cancer treatment protocols. Although they are not specific, some classical cancer drugs target centromeres and mitotic structures. These drugs may also destroy cells containing anaphase bridges so this mechanisms does not apply to every treated cancer.

BRCA1 and BRCA2 mutations increase the numbers of breaks, but breakage-fusion-bridge cycles also occur in sporadic breast cancer chromosomes, albeit less frequently. In addition to gene defects in BRCA1 and BRCA2, other inherited gene defects increase susceptibility to EBV infection and EBV-driven diseases. These inherited forms are associated with mutations in SH2D1A, ITK, MAGT1, CTPS1, CD27, CD70, CORO1A, and RASGRP1 (Latour and Winter, 2018). Over 50% of patients with one of these defects experience EBV-driven lymphoproliferative disease, including Hodgkin and non-Hodgkin lymphomas. Severe viral infections with other herpesviruses (CMV, HSV, HHV-6) are also common. EBV variants produce nucleases that can also cause breaks directly (Wu et al. 2010), promoting telomere dysfunction and replication stress (Hafez and Luftig, 2017). The idea that EBV variants initiate breaks is firmly established but that they also interfere with rejoining of fragments is somewhat speculative. This interference only needs to happen once according to the graphical abstract. Abundant EBV binding sites exist as homologous repetitive human DNA sequences nearly 2500 base pairs long. Other arguments also support this view. EBV DNA gets packaged around chromatin like a small human chromosome. The viral chromosome moves around through the nuclear environment of human chromosomes, depending on the state of viral activation. In its latent form, EBV binds human repressive heterochromatin, but reactivation moves the virus to active euchromatin (Moquin et al. 2018). DNA near breaks is likely to be accessible because helicases and other enzymes expose active chromatin (Aleksandrov et al. 2020). Exposure is necessary so that repair processes can access broken DNA.

Targeting mitosis in rapidly dividing cancer cells has been effective in traditional cancer therapy. Chemotherapy agents that disrupt microtubule function have been clinically valuable and include drugs such as taxanes and vinca alkaloids. These drugs interfere with microtubules in rapidly dividing cells and alter breakage fusion bridge formation. Managing accompanying side effects, toxicity and multi drug resistance is difficult (Ruan *et al*, 2018). Strategies to prevent cancer recurrence that target breakage-fusion-bridge formation more specifically may lead to drugs that destroy cells with two centromeres.

The current evidence supports cancer prevention by a childhood herpes vaccine and cancer treatment by EBV antivirals. The prospects for producing an EBV vaccine are promising, but the most appropriate targets are still not settled. Some immunotherapy strategies rely on augmenting the immune response, but this approach may need modification because mutations create additional holes in the immune response. Retroviruses and retrotransposons (Helman *et al*, 2014) may also participate in breast cancer breaks. Participation from porcine endogenous retroviruses is actionable by thoroughly cooking pork products. However, despite assertions that xenotransplantation with pig cells is safe, it is concerning that up to 6500 bps in human chromosome 11 are virtually identical to pig DNA.

It is unlikely that the extensive homology to EBV represents contamination of the reference human genome sequence. Most similarities occur to the unusual tumor viral variants HKHD40 and HKNPC60 and not EBV. Porcine Endogenous Retrovirus (PERV) or other endogenous retroviral sequences also contribute to numerous breast cancer breaks.

## Materials and Methods

### Breast cancer genomic sequences

#### Characteristics of hereditary breast cancers compared to viral cancers

Selection of hereditary breast cancer genomes for this study required patient samples with a known, typed *BRCA1* or *BRCA2*. gene mutation (Nik-Zainal *et al*, 2016; Nones *et al*, 2019). These hereditary cancers were mainly stage III ductal or no specific type (Nones *et al*, 2019). Seventy-four hereditary or likely hereditary breast cancers were selected as typed BRCA1 or BRCA2 mutation-associated cancers. Breast cancers diagnosed before age 40 did not have typed BRCA mutations, but they should be suspected in this group (Petrucelli *et al*, 1993). About half the women with breast cancer diagnosed before age 30 and with strong family histories of the disease had mutations in BRCA1, BRCA2 and TP53 (Lalloo *et al*, 2006). Although this group approximates BRCA1 and BRCA2 mutation carriers, its members were kept separate. A study of familial breast cancers contributed another 65 BRCA1/BRCA2 associated breast cancers (Nones *et al*, 2019). Results were checked against breakpoints in 101 triple-negative breast cancers from a population-based study (Staaf *et al*, 2019). Genome sequencing had been done before treatment began. Male breast cancers and cancers with BRCA1 or BRCA2 mutations diagnosed after age 49 were excluded since such mutations are less likely to be pathogenic. Cancers with hereditary mutations in PALB2 and p53 were also excluded. Sporadic breast cancers were taken as those diagnosed after age 70 without known inherited mutation.

#### Hereditary and sporadic breast cancer patient DNA sequence data

Sampling the population to represent the breadth of all somatic and hereditary breast cancers is not a trivial problem and limits many studies. There is no assurance that even large numbers of breast cancers are adequate because they are not a random sample from all breast cancers (Friedenson, 2009). The data used here comes from 560 breast cancer genome sequences, familial cancer data from 78 patients, methylation data from 1538 breast cancers vs. 244 controls, 243 triple-negative breast cancers and 2658 human cancers (Batra *et al*, 2021; Cortes-Ciriano *et al*, 2020; Nik-Zainal *et al*, 2016; Nones *et al*, 2019; Staaf *et al*, 2019). Gene breakpoints for inter-chromosomal and intra-chromosomal translocations were conveniently obtained from the COSMIC catalog of somatic mutations as curated from original publications or from original articles (Nik-Zainal *et al*, 2016; Nones *et al*, 2019; Staaf *et al*, 2019). NPC breakpoint positions were from Bruce et al. (Bruce *et al*, 2021). Burkitt’s lymphoma breakpoints were from Walker et al. (Walker *et al*, 2014).

#### Comparisons of DNA sequences

The GrCH38 human genome version was used whenever possible but some comparisons of chromothripsis breakpoints were easier with the GrCh37/ Hg19 version. piRNA locations were from the piRNA bank and converted to GrCh38 (Sai Lakshmi and Agrawal, 2008). The UCSC online web “Liftover” function interconverted different locations of genome coordinates such as those for viral-human similarities and piRNA locations. DNA flanking sequences at breakpoints were downloaded primarily from the UCSC genome browser as FASTA files and inputted directly into BLAST. Positions of differentially methylated regions near breast cancer breakpoints (Tang *et al*, 2012) were compared to breakpoint positions in a set of 70 NPCs based on data from Bruce et al. (Bruce *et al*, 2021). The nearest break position in breast cancer to a break in NPC was taken as the closest NPC breakpoint 5’ to the breast cancer break or the closest NPC value 3’ to the breast cancer break, whichever was closer. Vast difference in the amount of data available for NPC vs breast cancer breakpoints limited these comparisons near the ends of chromosomes. In these cases the minimum distance returned from the breast cancer break to the NPC break was either the position of the breast cancer break or the position of the NPC break. There were surprisingly few positions like this relative to the total numbers of breast cancer breaks so imposing this limit made little difference. The validity of the calculations was further tested for a few chromosomes by scanning the breast cancer breakpoints vs the NPC breakpoints in the reverse direction. The same results were obtained.

The NCBI BLAST program (MegaBLAST) and database (Altschul *et al*, 1990; Mount, 2007; Zhang *et al*, 2000) compared DNA sequences around breakpoints in breast cancers to all available viral DNA sequences. E(expect) values are related to p values and represent the probability that a given homology bit score occurs by chance. E values <1e-10 were considered to represent significant homology. In many cases, Expect values were “0” (<1e-180) and always far below 1e-10. Virus DNA was from BLAST searches using “viruses (taxid:10239)” with human sequences and uncharacterized sample mixtures excluded. Different strains and isolates of the same virus were tested for human homology as a group. Specifically, HKHD40 and HKNPC60 were considered together as “EBV”. Homology between these virus groupings vs. human sequences was determined using “Blastn”.

EBV DNA binding locations on human chromosomes were obtained from a publication (Lu *et al*, 2010) identifying genes within or near EBNA1 binding sites. Breaks in breast cancers were compared to EBV DNA binding sites.

#### Data analyses

Microsoft Excel, SPSS, StatsDirect, Visual basic, and Python + Biopython (Cock *et al*, 2009) scripts provided data analysis. The NCBI Genome Decoration page and the Ritchie lab website (Wolfe *et al*, 2013) provided chromosome annotation software.

#### Statistics

Statistical analyses used StatsDirect or SPSS statistical software. A scatter plot of the data was always prepared before running any statistical analysis. The Fisher exact test compared viral similarities around breakpoints in hereditary breast cancers to similarities in breakpoint positions generated by random numbers. Tests for normality included kurtosis and skewness values and evaluation by methods of Shapiro-Francia and Shapiro-Wilk (Shapiro, 1965). Details are included with the breakpoint calculations in Table S1.

#### Fragile site sequence data

Positions of fragile sites were from a database (Kumar *et al*, 2019) and original publications (Maccaroni *et al*, 2020). The presence of repetitive di- and trinucleotides tested for the exact positions of fragile sites. “RepeatAround” tested sequences surrounding breakpoints for 50 or fewer direct repeats, inverted repeats, mirror repeats, and complementary repeats.

## Supporting information

Brc vs NPC calculations

